# DNA 5-Hydroxymethylcytosines from Cell-free Circulating DNA as Diagnostic Biomarkers for Human Cancers

**DOI:** 10.1101/163204

**Authors:** Wenshuai Li, Xu Zhang, Xingyu Lu, Lei You, Yanqun Song, Zhongguang Luo, Jun Zhang, Ji Nie, Wanwei Zheng, Diannan Xu, Yaping Wang, Yuanqiang Dong, Shulin Yu, Jun Hong, Jianping Shi, Hankun Hao, Fen Luo, Luchun Hua, Peng Wang, Xiaoping Qian, Fang Yuan, Lianhuan Wei, Ming Cui, Taiping Zhang, Quan Liao, Menghua Dai, Ziwen Liu, Ge Chen, Katherine Meckel, Sarbani Adhikari, Guifang Jia, Marc B. Bissonnette, Xinxiang Zhang, Yupei Zhao, Wei Zhang, Chuan He, Jie Liu

## Abstract

DNA modifications such as 5-methylcytosines (5mC) and 5-hydroxymethylcytosines (5hmC) are epigenetic marks known to affect global gene expression in mammals(*1, 2*). Given their prevalence in the human genome, close correlation with gene expression, and high chemical stability, these DNA epigenetic marks could serve as ideal biomarkers for cancer diagnosis. Taking advantage of a highly sensitive and selective chemical labeling technology(*3*), we report here genome-wide 5hmC profiling in circulating cell-free DNA (cfDNA) and in genomic DNA of paired tumor/adjacent tissues collected from a cohort of 90 healthy individuals and 260 patients recently diagnosed with colorectal, gastric, pancreatic, liver, or thyroid cancer. 5hmC was mainly distributed in transcriptionally active regions coincident with open chromatin and permissive histone modifications. Robust cancer-associated 5hmC signatures in cfDNA were identified with specificity for different cancers. 5hmC-based biomarkers of circulating cfDNA demonstrated highly accurate predictive value for patients with colorectal and gastric cancers versus healthy controls, superior to conventional biomarkers, and comparable to 5hmC biomarkers from tissue biopsies. This new strategy could lead to the development of effective blood-based, minimally-invasive cancer diagnosis and prognosis approaches.

## INTRODUCTION

Cytosine methylation (5mC) is a well-established epigenetic mechanism that affects global gene expression. It is extensively remodeled during mammalian development and cell differentiation, as well as during cancer initiation, progression, and therapeutic response(*4, 5*). Active demethylation in the mammalian genome is mediated by the TET family of dioxygenases 2 that oxidize the 5mC modification to 5-hydroxymethylcytosine (5hmC)(*6, 7*), and further to 5- formylcytosine (5fC), and 5-carboxylcytosine (5caC)(*8-10*). The “intermediate” 5hmC not only marks active demethylation but also serves as a relatively stable DNA mark that plays distinct epigenetic roles(*2, 11-16*). Recent genome-wide sequencing maps of 5hmC in various mammalian cells and tissues support its role as a marker for gene expression(*17-23*); it is enriched in enhancers, gene-bodies, and promoters, and changes in 5hmC correlate with changes in gene expression levels(*23, 24*).

The discovery of cell-free DNA (cfDNA) originating from different tissues in the circulating blood has revolutionary potential for the clinic(*25*). Liquid biopsy-based biomarkers and detection tools offer substantial advantages over existing diagnostic and prognostic methods, including being minimally invasive, which will promote higher patient compliance, clinically convenient, cost-efficient, and enabling dynamic monitoring(*26*). Tumor-related somatic mutations in cfDNA have been shown to be consistent with the tumor tissue, although low mutation frequency and the lack of information on tissue of origin hamper the detection sensitivity. 5mC and 5hmC in cfDNA from liquid biopsies could serve as parallel or more valuable biomarkers for non-invasive diagnosis and prognosis of human diseases because they recapitulate gene expression changes in relevant cell states. If these cytosine modification patterns can be sensitively detected, disease-specific biomarkers could be identified for effective early detection, diagnosis and prognosis.

High-throughput sequencing is an ideal platform for detecting genome-wide cytosine modification patterns. Whole genome bisulfite sequencing or alternative reduced representative methods have been applied in biomarker research with cell-free DNA(*27-29*). Tissue- and cancer-specific methylation sites have shown promising performance in tracking tissue-of-origin3 from circulating blood(*27, 29*). However, 5mC serves mostly as a repressive mark with a high background level in the human genome, and its sequencing with bisulfite treatment has been hampered with extensive DNA degradation, in particular with cfDNA. Taking advantage of the presence of the hydroxymethyl group, selective chemical labeling can be applied to map 5hmC using low-input DNA with high sensitivity. The profiling method is robust and cost-effective for large cohort studies and practical applications. Here, we established the 5hmC-Seal technology for 5hmC profiling in cell-free DNA. We show that the differentially enriched 5hmC regions in cfDNA are excellent markers for solid tumors.

## RESULTS

### Overview of the nano-hmC-seal profiling in clinical specimens

We optimized our previously published profiling method(*3*) (Fig. 1) for cell-free DNA. The adaptor is pre-ligated with barcodes to enhance the library construction efficiency and decrease the cross contamination between large cohort of samples. The labeling, binding and washing steps are optimized for capturing limited 5hmC-containing cfDNA fragments. We profiled 5hmC in plasma cfDNA from cancer patients and healthy controls, as well as in genomic DNA (gDNA) isolated from tumors and adjacent healthy tissue including 90 healthy individuals, 260 cancer patients, and 71 patients with benign diseases among Chinese populations (Table S1, S2). For these patients and healthy controls, the study generated 401 hmC-Seal libraries from plasma cfDNA and 188 hmC-Seal libraries from tissue gDNA (Table S3). The cohort samples were collected and profiled in three batches (Table S3). To minimize the influence of experimental batch effect, differential 5hmC between cancers and controls (or between tumors and adjacent tissues) were analyzed with the first (discovery) batch and validated in the second (validation) and third (additional validation) batches.

**Figure 1.**
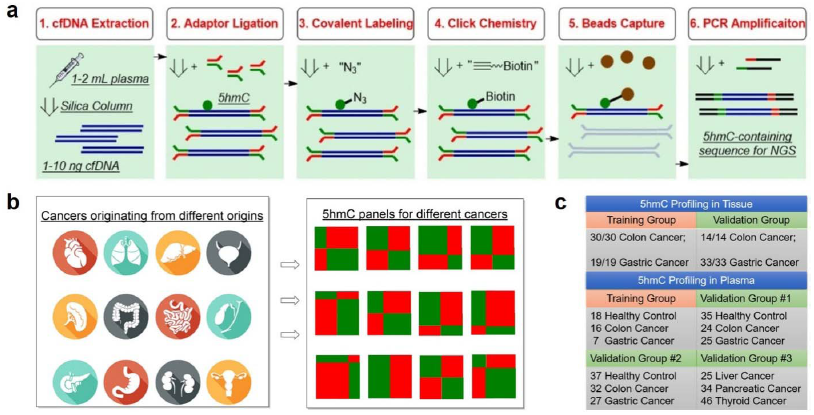
Detecting 5hmC biomarkers in cfDNA of human cancers. **a**, Workflow of 5hmC-Seal profiling from cfDNA is shown. Purified cfDNA is ligated with standard sequencing adaptors. 5hmC-containing cfDNA fragments are selectively labeled with a biotin group. The biotin-labeled fragments are captured on the avidin beads, followed by PCR amplification and next-generation sequencing (NGS). **b**, Cancers of different origins may release cfDNA decorated with distinct 5hmC modification patterns. Unique 5hmC signatures specific for different cancer types could be detected as biomarkers for diagnosis and prognosis. **c**, Schematic overview of sample collection, data generation and analysis.

To validate the 5hmC capture efficiency and reliability of the modified assay, we spiked a pair of synthesized 5hmC-containing and non-5hmC-containing DNA probes into plasma cfDNA. The 5hmC-Seal capture generated an average of 56-fold 5hmC enrichment of the spike-in probes compared to control without pull-down (Fig. S1a). Samples with physiologically relevant amounts of cfDNA (1, 2, 5, 10, 20 ng) and spike-in 5hmC-containing probes were processed and sequenced, respectively. A linear relationship was observed between the proportion of 5hmC-containing spike-in readouts and the spike-in concentration within cfDNA (*r*^2^=0.99, Fig. S1b), confirming a quantitative 5hmC capture even down to 1 ng of input cfDNA.

### Global and genomic distribution of 5hmC modifications

We evaluated the global 5hmC level variation in cancer by using an ultra-sensitive capillary electrophoresis-electrospray ionization-mass spectrometry (CE-ESI-MS) method(*30*). Global 5hmC levels of the tumor gDNA markedly decreased compared to the adjacent healthy tissue gDNA, with an average of 85% and 64% reduction in colorectal and gastric tumor, respectively. The global 5hmC levels of the cancer plasma cfDNA showed a more limited decrease compared to control plasma cfDNA, consistent with low proportions of tumor-derived DNA in the total cfDNA pool (Fig. S2).

In plasma cfDNA, 5hmC is enriched within gene bodies and DNase I sensitive peaks, while depleted at transcription start sites, CpG islands and transcription factor (TF) binding peaks relative to the flanking areas (Fig. S3a-f), suggesting accumulation of 5hmC surrounding TFs at active transcription sites. 5hmC is also enriched in several permissive histone marks such as H3K27ac, H3K4me1 and H3K9me1, while repressive markers such as H3K9me3 are underrepresented (Fig. S3g-r). The genomic distribution of 5hmC in tissues gDNA is generally consistent with that observed in plasma cfDNA samples (Fig. S3). The median distribution of 5hmC is also similar between disease and healthy samples (Fig. S4).

### Differential 5hmC loci associated with colorectal cancer

The average 5hmC profile of plasma cfDNA is distinct from that of tissue gDNA (Fig. 2a), which could be due to their distinct cell origins and/or the different DNA degradation properties in cell free circulation. Variations attributable to tissue identity (cell free plasma, white blood cells, colon and stomach tissues) are dominant over variations attributable to disease status (healthy individual versus cancer patient, tumor versus adjacent tissue). In addition, plasma cfDNA of colorectal and gastric cancers are more closely related with each other than with plasma cfDNA of healthy controls (Fig. 2a), implicating common variations in different cancer types.

**Figure 2.**
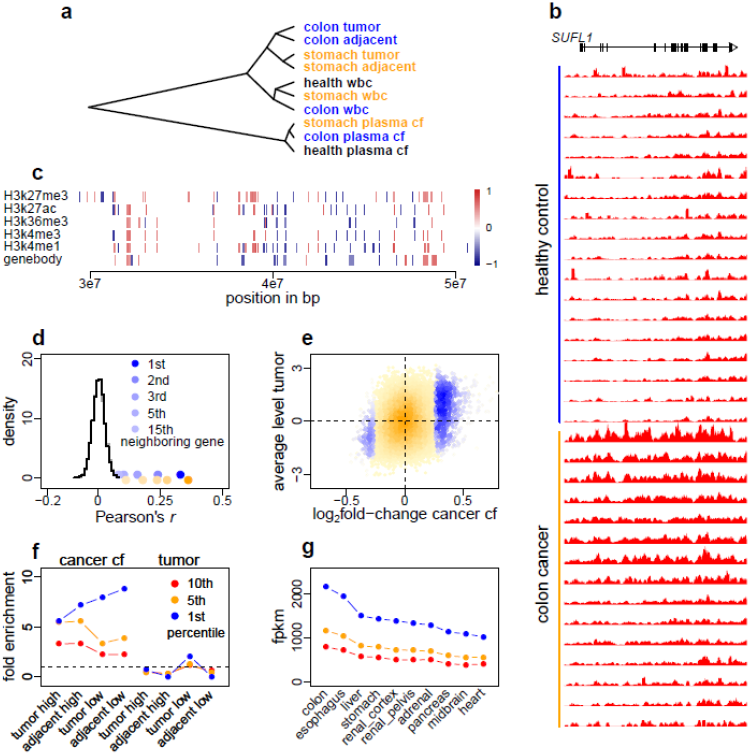
Differential 5hmC loci associated with cancer. **a**, Average 5hmC levels in gene body across healthy controls (health) and cancer patients (colon and stomach), estimated for plasma cfDNA (plasma cf), white blood cell genomic DNA (wbc) and tissue genomic DNA (tumor and adjacent), were clustered by correlation distance. **b**, Counts per million reads at *SULF1* gene (plus +/-20kb region) in plasma cfDNA of the 15 healthy controls and 18 colorectal cancer patients. The moving averages at 0.01 smoother span are shown. **c**, The distribution of colorectal cancer-associated 5hmC loci detected at 5% false discovery rate in plasma cfDNA. The color key indicates relative change. **d**, Pearson’s correlation of log_2_ fold changes between analyzed genes and their neighboring genes was plotted against the null distribution of correlation between genes and their 1st neighboring genes, generated by shuffling gene positions for 1000 times. Blue and orange points denote data from plasma cfDNA and tissue gDNA, respectively, for colorectal cancer. In **c** and **d** chromosome 1 are shown for an example. **e**, The average 5hmC levels in gene bodies in tumor gDNA were plotted against the log_2_ fold change of 5hmC levels in colorectal cancer plasma cfDNA. Orange points denote analyzed genes and blue points denote differential genes called at 5% FDR and 1.2 fold change, color intensity representing data density. **f**, Enrichment of genes with cancer-associated 5hmC level increase (or decrease) in genes with high (or low) 5hmC levels in tissues (tumor and adjacent). The 1st, 5th and 10th percentile genes in descending or ascending order of the log_2_ fold change were compared against the corresponding percentile genes in descending or ascending order of the average 5hmC levels. Cancer cf: differential genes detected in cancer plasma cfDNA; tumor: differential genes detected in tumor tissue. Dashed line denotes no enrichment. **g**, 5hmC reads from plasma cfDNA of 15 colorectal cancer patients were summed over ENCODE DNase broad peaks derived from various tissues. Tissues were ordered by the 1st percentile of fragments per kilo-bases per million (fpkm) in descending order. For each tissue, 20000 DNase peaks were randomly sampled from one ENCODE tissue sample with good sequencing quality.

We compared 5hmC profiles from plasma cfDNA between 15 colon cancer patients and 18 healthy controls in the discovery batch to identify differential 5hmC loci. The profiles were separated into 18 feature categories: gene bodies, promoters, CpG islands, and *cis*-regulatory elements delineated by the Encyclopedia of DNA Elements (ENCODE)(*31*). A parallel analysis compared 5hmC profiles from gDNA between colorectal tumors and adjacent tissues in 30 patients in the tissue discovery batch. All feature categories showed enrichment of differential 5hmC loci (Table S4). Fig. 2b shows a differential locus detected in plasma cfDNA at the *SULF1* (sulfatase 1) gene. In cancer plasma cfDNA, the 5hmC levels in *SULF1* are elevated in both exons and introns, with a peak pattern similar to that of tissue gDNA (Fig. S5a). Differential 5hmC loci across feature categories, particularly gene bodies and histone modification peaks, show regionally elevated or decreased 5hmC levels along neighboring loci (Fig. 2c). Indeed, correlation of cancer-associated 5hmC changes between neighboring genes is significantly higher than a null distribution generated by shuffling gene positions within chromosome (Fig. 2d). This may suggest that 5hmC modifications occur and change in a relatively long-range, region-wise pattern.

Across the genomic features, the average 5hmC levels normalized by feature length are more or less correlated between the cancer plasma cfDNA and tumor gDNA samples (Spearman’s ρ 0.34-0.84, Fig. S5b). This locus-specific correlation of the 5hmC level is expected because of biological constraints. In contrast, we found no correlation between the log_2_ fold change of 5hmC levels in cancer plasma cfDNA and that in tumor gDNA (Fig. S5b). Because 5hmC levels show greater variations among different tissues than between disease status (Fig. 2a), when gDNA from tumor tissue is released into plasma and mixed with the vast amount of background cfDNA derived from a variety of different tissues, the additional tumor signal observed at a given locus would be determined by the order of locus-, tissue- and disease-specific variations. Consistent with this expectation, we found that genes with 5hmC level elevated in cancer patient’s plasma cfDNA were enriched in genes with high 5hmC levels in the tumor tissue gDNA (Fig. 2e). Specifically, the top 1% genes with 5hmC level most elevated in cancer plasma cfDNA were enriched by over five-fold in the top 1% genes with greatest 5hmC levels in tumor and adjacent tissues (Fisher’s exact tests P <0.001, Fig. 2f). Similarly, genes with 5hmC level decreased in cancer plasma cfDNA were enriched in genes with low 5hmC levels in tumor and adjacent tissues (Fig. 2e, 2f). In contrast, no such enrichment pattern was observed for the differential 5hmC loci detected in tumor gDNA (Fig. 2f). To further investigate the tissue origin of cancer plasma cfDNA, 5hmC reads from the 15 colorectal cancer patients were summed over ENCODE DNase hypersensitivity peaks derived from various tissues of healthy individuals. The peaks derived from colon tissue contain the greatest amount of 5hmC modifications compared to the peaks derived from other tissues (Fig. 2g), indicating the tissue specificity of 5hmC signals in cancer plasma cfDNA.

### Classification of colorectal cancer by 5hmC markers derived from plasma cfDNA

Unsupervised hierarchical clustering using differential 5hmC loci derived from plasma cfDNA generally separated colorectal cancer patients from healthy individuals in the validation batch (Fig. 3a). Across the various feature categories, the log_2_ fold change of 5hmC level in gene bodies showed greatest correlation between the discovery and validation batches (Spearman’s ρ=0.79, Fig. 3b), indicating that 5hmC loci in gene bodies are potentially more stable cancer biomarkers. We selected 989 differential loci in gene bodies detected at 5% false discovery rate (FDR) and 1.2-fold change (increase or decrease in cancer; Table S5) for cancer classification. Specifically, a model-based classifier that applies elastic net regularization on logistic regression was trained using the discovery samples (15 patients vs. 18 controls) and then tested in the validation samples (24 patients vs. 35 controls). Receiver operating characteristic (ROC) curves were generated to evaluate the performance using the area under the ROC curve (AUC). The prediction algorithm achieved 83% sensitivity and 94% specificity (AUC = 0.95, Fig. 3c) for patient classification.

**Figure 3.**
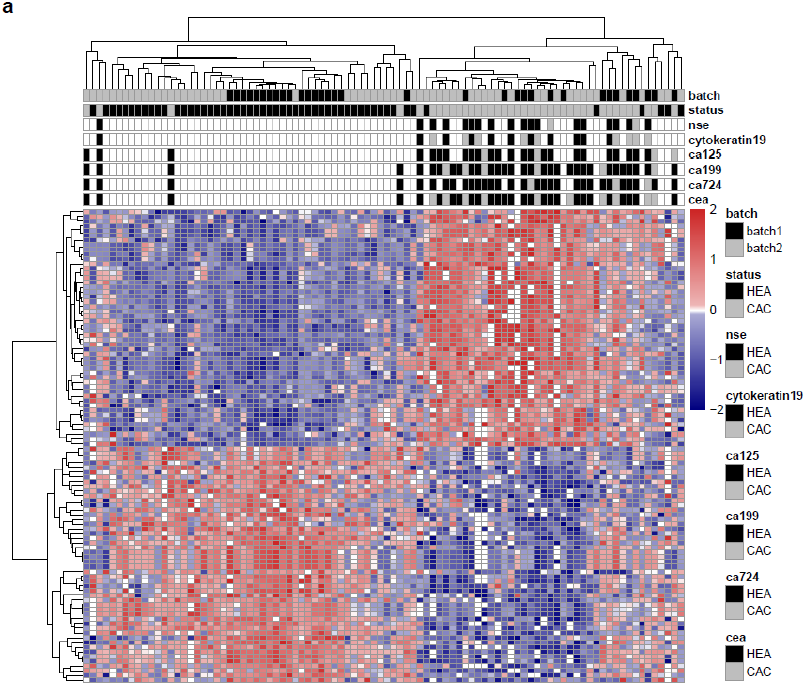
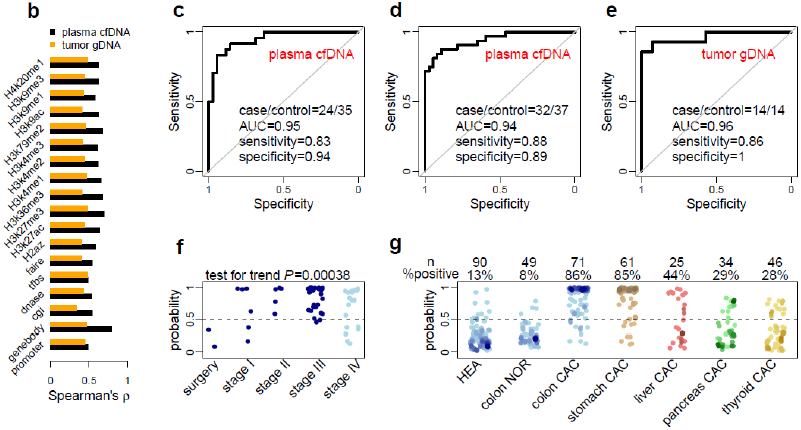
Performance of 5hmC biomarkers for colorectal cancer patients. **a**, The heatmap shows clustering using the 50 most up-regulated and 50 most down-regulated 5hmC loci from the discovery batch across cfDNA samples from both the discovery and validation batches. Diagnostic results using classical biomarkers are also shown. HEA: healthy individuals; CAC: cancer patients; nse: neuron specific enolase; CEA: carcinoembryonic antigen; CA125/19-9/72-4: carbohydrate antigen 125/19-9/72-4. **b**, Correlation of 5hmC variation in cancer between the discovery and validation batches of samples is higher in plasma cfDNA (cancer patients vs. healthy individuals) than in tissue genomic DNA (tumors vs. adjacent tissues), especially for 5hmC loci in gene bodies. **c-d**, Classifying two independent validation batches using 5hmC classifier derived from plasma cfDNA from the discovery batch. **e**, Classifying an independent set of colon cancer tumor tissues using 5hmC biomarkers detected from the discovery batch of tissue samples (tumors vs. adjacent tissues). AUC: area under curve. **f**, The predicted probability (score) of a cancer patient based on 5hmC biomarkers from plasma cfDNA shows a significant trend associated with clinical stage. Patients after surgery show predicted scores undistinguishable from healthy individuals. **g**, The 5hmC biomarkers detected in plasma cfDNA from colorectal cancer patients are potentially cancer type-specific, showing differential predictive performance in plasma cfDNA from stomach, liver, pancreatic and thyroid cancer patients. HEA: healthy control; NOR: patient with benign tumor; CAC: cancer patient.

An additional validation batch (32 patients vs. 37 controls) was independently collected and tested, achieving 88% sensitivity and 89% specificity (AUC = 0.94, Fig. 3d). The classifier was further tested on a set of U.S. samples (Table S2), all of European descent, collected at the University of Chicago Medical Center. The classifier detected 4 out of 5 patients as cancer positive (80% sensitivity) and called 1 out of 6 healthy controls (83% specificity) in this small cohort. Therefore, although the classifier was trained on Chinese patients, it could capture a general signal of 5hmC changes in plasma cfDNA in colorectal cancer.

For comparison, we applied similar approaches to evaluate the performance of 5hmC biomarkers derived from colorectal tumor tissues (Table S1). We selected 219 differential loci at gene bodies called at 5% FDR and 1.2-fold change between tumor and adjacent tissues (Table S6) from 30 patients of the discovery batch. The 5hmC tissue biomarkers showed a sensitivity of 86% and a specificity of 100% (AUC = 0.96, Figure 3e) in 14 patients from the tissue validation batch, suggesting that the 5hmC biomarkers from plasma cfDNA exhibits performance comparable to that from tissue gDNA.

### Disease sensitivity and specificity of plasma cfDNA-derived 5hmC markers

We next assessed the ability of the 5hmC biomarkers derived from plasma cfDNA to classify cancer stages in patients with available records. The 5hmC classifier assigned incremental numbers of cancer individuals (predicted cancer probability > 0.5) for patients with surgery treatment (0/2), patients at cancer stage I (4/6), and patients at stage II and III (38/40) (Cochran–Armitage test for trend P = 0.00038, Fig. 3f). The classifier had good but reduced power to call Stage IV patients (12/18), who in general suffered from metastasis to various tissues and are expected to have more complex tumor DNA profiles in circulation due to metastasis.

We further assessed disease and tissue specificity of the classifier in patients with colon-related benign diseases (n=49) and patients with colorectal (n=71), gastric (n=61), liver (n=25), pancreas (n=34) and thyroid (n=46) cancer. Compared to the 86% call rate in colorectal cancer patients, only 8% of patients with benign colon diseases were predicted as cancer (Fig. 3g). The classifier also demonstrated certain tissue specificity, with a decreasing cancer calling rate in gastric (85%), liver (44%), pancreatic (29%) and thyroid (28%) cancer patients (Fig. 3g). The lower sensitivity in calling the other cancers is not due to intrinsic difficulty in distinguishing those cancers, as we achieved much greater sensitivity in liver and pancreatic cancer using 5hmC markers derived from plasma cfDNA of liver and pancreatic cancer patients respectively (data not shown). These results indicate that distantly related cancers can be readily distinguished through joint testing by the corresponding classifiers, while classification of closely related cancers such as colorectal and gastric cancer may be facilitated by additional diagnostic criteria.

A subset of cancer patients had records of classical biomarkers and epidemic factors, with which we compared plasma cfDNA-derived 5hmC biomarkers for cancer detection sensitivity. The detection sensitivity of carcinoembryonic antigen (CEA, 32%), alpha-fetoprotein (AFP, 0%), carbohydrate antigen 125 (CA125, 13%), CA15-3 (0%), CA19-9 (19%), CA72-4 (17%), cytokeratin 19 (49%), Neuron-Specific Enolase (NSE, 21%), overweight (body mass index ≥25 kg/m^2^, 34%), smoking (9%), alcohol consumption (7%) and previous history of cancer (0%) were all less than 50%. By calling cancer if any conventional biomarker or risk factor is positive, the upper bound detection sensitivity of the combined classical biomarkers and epidemic factors only reached 54%, a sensitivity rate much lower than the 86% that we could achieve using 5hmC markers. In addition, compared with the methylated *SEPT9* gene (encoding septin 9), a blood-based epigenetic biomarker for colon cancer, our cfDNA 5hmC biomarkers registered a significantly further improved overall sensitivity (0.86 versus 0.48 based on public data)(*32*).

### 5hmC markers derived from plasma cfDNA in gastric cancer

Next, we analyzed gastric cancer using plasma cfDNA samples. In the discovery batch, 5hmC loci in 7 gastric cancer patients were compared to 18 healthy controls across genomic features (Table S4). Using the top 100 elevated or decreased 5hmC loci, 25 gastric cancer patients could be generally separated from 35 healthy individuals in the validation batch (Fig. S6a). Again, 5hmC changes in gene bodies showed relatively higher correlation between the discovery and validation batches compared to other genomic features (Fig. S6b). A model-based classifier was generated using the 1,431 differential loci in gene bodies identified at 5% FDR and 1.2-fold change in the discovery batch (Table S7), and was applied to the validation batch, achieving 92% sensitivity and 91% specificity (AUC = 0.93, Fig. S6c). Further assessment of the gastric cancer classifier in an additional validation batch collected independently (29 patients vs. 37 controls) achieved 90% sensitivity and 97% specificity (AUC = 0.97, Fig. S6d). The classification performance of the 5hmC biomarkers derived from cancer cfDNA was comparable to that from tumor gDNA samples as well: 161 differential 5hmC loci in gene bodies detected in 19 pairs of tumors and adjacent tissues in the discovery batch (Table S8) was applied on 33 pairs of tissues in the validation batch, achieving 82% sensitivity and 94% specificity (AUC = 0.93, Fig. S6e).

The 5hmC gastric cancer classifier derived from plasma cfDNA showed a trend of increasing cancer calling (predicted cancer probability > 0.5) rate with cancer severity (P = 0.11, Fig. S6f). The classifier also demonstrated disease and tissue specificity, with 0% cancer calling rate for benign gastric diseases, and with decreasing cancer call rate in patients with colorectal (61%), liver (28%), pancreatic (6%) and thyroid (0%) cancer patients (Fig. S6g).

The detection sensitivity of classical biomarkers and epidemic factors for gastric cancer was 13% (CEA), 6% (AFP), 6% (CA125), 3% (CA15-3), 14% (CA19-9), 29% (CA72-4), 36% (cytokeratin 19), 13% (NSE), 18% (overweight), 25% (smoking), 10% (alcohol) and 7% (previous history of cancer). The upper bound sensitivity combining these markers and factors was 70%, which again is lower than the 80% sensitivity observed in our studies.

### Tissue origin of the cancer associated 5hmC changes observed in plasma cfDNA

To demonstrate the tumor relevance of plasma cfDNA, we sought to examine its source in patient-derived xenograft (PDX) mouse models. PDX mouse models were derived from tumors of three colorectal patients and three gastric patients, each with three independent xenograft animals. Plasma cfDNA of PDX mice was collected at 12-15 weeks of age, from which the 5hmC-containing fragments were enriched and sequenced using the same protocol as with human plasma cfDNA. The proportion of cfDNA derived from the tumor, estimated as the proportion of sequencing reads uniquely mapped to the human genome, was significantly increased in mice grafted with gastric tumors (P = 0.0020) and showed a trend of increase in mice grafted with colorectal tumors with fewer passages (P = 0.16, Fig. 4a). Only the sequencing reads mapped to human genome were further analyzed.

**Figure 4.**
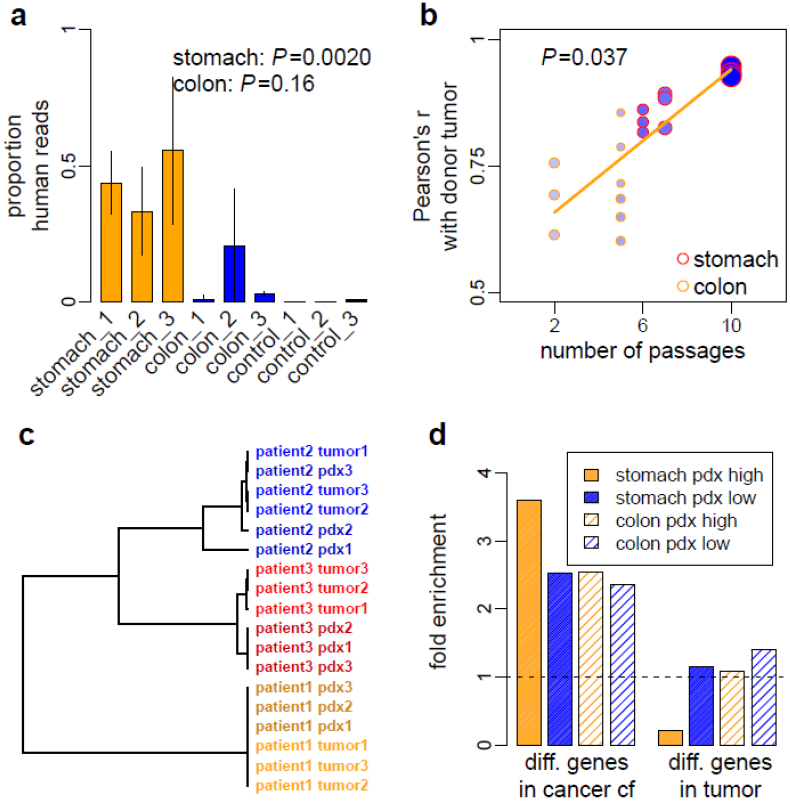
The origin of cancer associated 5hmC changes observed in plasma cfDNA. **a**, The proportions of human reads in plasma cfDNA captured with hmC-Seal are shown for PDX mice grafted with tumor from three gastric cancer patients (stomach_1 to 3), three colorectal cancer patients (colon_1 to 3) and for PDX mice without graft (control_1 to 3). Vertical bars represent standard deviation estimated from three replicate PDX mice for each patient. The PDX mice grafted with gastric tumor had greater number of passages (6-10) than those grafted with colorectal tumor (2-5). **b**, The correlation of the 5hmC profile between tumor-derived, PDX plasma cfDNA and donor tumor gDNA depends on the number of passages of the PDX mouse. The size of the points is proportional to the size of grafted tumor and the density of color denotes the growth rate of the grafted tumor. **c**, Using the correlation distance of the top five genes that had the greatest 5hmC level in PDX plasma cfDNA, donor tumor gDNA and PDX plasma cfDNA from the same individual patient were clustered together. **d**, Directional enrichment of genes with altered 5hmC levels in cancer plasma cfDNA or in tumor tissue gDNA in genes with high or low 5hmC levels in PDX plasma cfDNA. The top 5% genes in descending or ascending order of the log_2_ fold change were compared against the corresponding top 5% genes in descending or ascending order of the average 5hmC levels across three PDX replicates, derived from one gastric cancer patient (10 passages) and one colorectal cancer patient (5 passages). Dashed line denotes no enrichment.

Pearsonc’s correlation of 5hmC profile between plasma cfDNA of PDX mice and gDNA of donor tumors significantly depends on the number of passages (P = 0.037, Fig. 4b). This suggests a quantitative relationship between tumor growth and the experimental capturing of tumor 5hmC in plasma cfDNA, as the size (P=0.0096) and growth rate (P=0.0080) of tumors grafted in PDX mice increase with passage numbers (Fig. 4b). Using the top five genes with the greatest 5hmC levels in PDX plasma cfDNA, donor tumor and the derived PDX from the same individual patient were clustered together (Fig. 4c), supporting donor tumor tissue as the origin of the PDX cfDNA.

PDX allowed us to study tumor-derived cfDNA without confounding from background cfDNA. Genes with greater 5hmC levels in tumor-sourced PDX plasma cfDNA are more likely to be the genes with elevated 5hmC level in plasma cfDNA in cancer patients. Indeed, we found that genes with 5hmC level increased in patient plasma cfDNA were enriched in those genes with greater 5hmC levels in PDX plasma cfDNA, whereas genes with 5hmC level decreased in patient plasma cfDNA were enriched in genes with lower 5hmC levels in PDX plasma cfDNA (Fisher’s exact tests P <1×10^−9^, Fig. 4d). In contrast, genes with 5hmC level changed between tumor and adjacent tissues showed no such enrichment pattern (Fig. 4d).

### Tumor-associated 5hmC changes in gene regulation

To investigate the potential functional role of 5hmC in gene regulation, we evaluated the relationship between gene expression changes and 5hmC level changes in tumors in two colorectal and one gastric cancer patients. We performed the RNA-seq assay in tumor tissues and paired adjacent tissues. The log_2_ fold changes of gene expression and the log_2_ fold change of 5hmC level in tumors relative to adjacent tissues were estimated across the three patients. Gene dysregulation and 5hmC changes were then compared across a combined list of 200 differential 5hmC loci in gene bodies detected in colorectal and gastric tumors in the discovery batches. The correlation between gene expression changes and 5hmC changes in tumors is highly significant (P = 9.8×10^−6^, Fig. S7a). In addition, genes with altered 5hmC level in cancer plasma cfDNA or in tumor gDNA were enriched in cancer- and metastasis-related pathways(*33*) (Fig. S7b).

## DISCUSSION

5hmC in general marks active loci, as gene activation requires removal of the repressive 5mC methylations. It occurs in gene bodies of activated genes as well as various enhancers, indicating that the genomic locations of 5hmC reflect both gene activation and active chromatin state. 5hmC is chemical stable and thus its locations in gDNA can be stored in fragmented cfDNA for potential non-invasive detections. Utilizing a robust and highly efficient profiling-based approach to map 5hmC in plasma cfDNA samples from patients with cancer, we were able to identify 5hmC biomarkers that can distinguish cancer patients from healthy individuals with high sensitivity and specificity for colorectal (Fig. 3) and gastric (Fig. S6) cancers. Our study revealed genome-wide patterns of cancer-associated 5hmC changes in plasma cfDNA (Fig. 2) and demonstrated the cancer relevance of cfDNA using PDX models (Fig. 4). We also found a strong co-localization between 5hmC and open chromatin at active transcription (Fig. S2) and confirmed a noticeable correlation between cancer-associated 5hmC changes and gene expression changes (Fig. S7), further supporting 5hmC as a marker for open chromatin state and gene activation. The identified 5hmC biomarkers can be cancer type-specific (Fig. 3, Fig. S6): optimization of this approach with additional patient studies in the future will further improve performance and expand the application scope to other cancers. The strategy presented here provides the foundation for effective future liquid biopsy-based diagnosis and potentially prognosis of human diseases using cfDNA.

The dynamics of cancer cfDNA turnover in circulation is yet largely unknown. Under a likely simplified model, gDNA of tumor tissue is released into plasma and undergoes degradation with an equilibrium similar to that of background cfDNA from normal healthy tissues. Locus-specific 5hmC modification appears to be the primary determinant of 5hmC variations, with tissue specificity and then cancer state adding additional layers of variations. These tissue-, and to a lesser extent, cancer-specific signals released from tumor tissues slightly shift the 5hmC modification profile of background plasma cfDNA toward that of tumor tissue gDNA. The more cfDNA released from tumor tissues the greater of the shift and the power to discriminate biological variation of the source tumor. Therefore, integration of a panel of 5hmC profiles from gDNA of diverse tissue types is critical for future assessment of disease specificity in cancer biomarkers. In addition, solid tumors are composed of carcinoma stem cells and carcinoma cells, with a microenvironment constituted by leucocytes, cells of mesenchymal origin, and extracellular matrix (*34*). Tumor progression initiates a gradient change in local environment characterized by hypoxia and vascularization. Extensive variability may exist within a growing tumor and its surrounding cells such that certain types of cells are prone to apoptosis and some are more prone to release DNA to circulation. We expect that cancer-associated 5hmC changes observed in plasma cfDNA were contributed by distinct sets of cells within or surrounding tumor tissues. Single-cell or cell-type-specific 5hmC profiling, by decomposing tumor tissues and using appropriate cell type markers, would reveal the extent and distribution of cell specificity and shed further light on the properties of the source cells that contribute to the cancer-associated 5hmC changes observed in plasma cfDNA. These are future directions we wish to pursue.

## MATERIALS AND METHODS

### Study Design

#### Patient population

A total of 180 patients older than 18 years with colorectal, gastric cancers and hepatocellular carcinoma were diagnosed at 3 different medical centers in Shanghai Huashan Hospital at Fudan University, China from September 2015 to July 2016. 80 patients with thyroid cancer and pancreatic cancer were diagnosed at Peking Union Medical College Hospital, China during 2014-2016. The population was socioeconomically diverse, most of whom came from Beijing and East China (City of Shanghai, Zhejiang Province, Jiangsu Province and Anhui Province). All were collected from patients who were newly diagnosed or with postoperative recurrence, and 2 patients were postoperative colorectal cancer patients, with confirmation by histological evaluation. Patients treated with chemotherapy, radiation therapy, or immunotherapy, were excluded from this study. In total, this retrospective cohort study was conducted among 80 colorectal cancer patients and 75 gastric cancer patients, with an additional 25 hepatocellular carcinoma patients, 34 pancreatic cancer patients and 46 thyroid cancer patients to explore cancer type-specificity. Whole blood samples from 90 healthy individuals under physical examination were also collected at Fudan University, China during September 2015-May 2016 as healthy controls; these individuals are Chinese and showed no history of cancer and had no abnormalities in laboratory examinations. However, the follow-up data for all patients were unavailable because of the short follow-up time. Informed consent was obtained from each participating subject before the study, which was approved by the Institutional Review Board at each collaborating institution.

#### Batch design

To minimize the influence of batch effect, gastrointestinal participants were assigned into 3 batches according to chronological order. Differential 5hmC between cancer and control was analyzed with batch 1 (discovery) and validated in batch 2,3 (validation and additional validation).

#### Sample overview

Detailed information of the study subjects is shown in Table S1, S2, including number of samples, gender, age, clinical diagnosis, stage classified according to the Tumor, Node and Metastasis (TNM) guidelines (version 7), and eight conventional cancer biomarkers were measured in patients. The average age of 80 GC patients and 75 CC patients were similar, with more male than female cancer patients. 84.4% percent (65 out of 75) of CC and 70.4% percent (50 out of 71) of GC were mainly at advanced stages (stage III and IV). The pathological grade of 74.6% (51 out of 66) CC patients was moderate, while 73.4% (47 out of 64) GC patients was poor differentiation. And 53.8% (35 out of 65) of CC and 62.9% (39 out of 62) of GC patients showed lymph node metastasis. 26.3% (20 out of 76) of CC and 18.3% (13 out of 71) of GC patients showed distal metastasis. Four common tumor markers for gastrointestinal cancer screening are CEA, AFP, CA19-9 and CA72-4, which were positive in 0, 33.8%, 21.4%, 16.9% of CC and 7.5%, 13.6%, 14.3%, 29.0% of GC patients respectively.

#### Additional U.S. samples for validation

Five patients with colorectal cancer were diagnosed at the University of Chicago Medical Center from September 2015 to July 2016. All samples were collected from patients who were newly diagnosed and had no distal metastasis at the time of blood draw. Whole blood samples from 6 healthy individuals under physical examination were also collected at the University of Chicago Medical Center during September 2015-May 2016 as healthy controls; these individuals are non-Hispanic or Latino individuals of European ancestry. Informed consent was obtained from each participating subject before the study, which was approved by the Institutional Review Board at the University of Chicago.

#### Preparation of cfDNA Samples

cfDNA samples were prepared from peripheral blood collected from patients and healthy controls. Briefly, 4ml of peripheral blood was collected from each subject using EDTA anticoagulant tubes, and the plasma sample was prepared within 6h by centrifuging twice at 1,350g for 12min, and then centrifuging at 13,500g for 12min. The prepared plasma samples (about 2ml/subject) were immediately stored at -80°C. The plasma cfDNA was isolated using the QIAamp Circulating Nucleic Acid Kit (Qiagen) according to the manufacturer’s protocol. Within each experimental batch, samples were randomized on disease status in the following library preparation and sequencing profiling.

#### Isolation of Genomic DNA from Tissues

Tissue samples, including tumor and adjacent tissue samples, from patients were stocked at -80°C after surgical operation. 10-25 mg tissue was collected using a scalpel after sample unfreezing. Genomic DNA from tissues was isolated using the ZR Genomic DNA-Tissue Kits (Zymo Research) according to the manufacturer’s protocol.

#### 5hmC-Seal-seq Library Preparation and Sequencing

Seal-seq libraries for 5hmC profiling were prepared following our previously patented technology(*3*). In this method, the T4 bacteriophage β-glucosyltransferase is used to transfer an engineered glucose moiety containing an azide group onto the hydroxyl group of 5hmC across the human genome. The azide group is then chemically modified with biotin for affinity enrichment of 5hmC-containing DNA fragments. First, the genomic DNA is fragmented using an enzymatic reaction. Next, the fragmented genomic DNA or the cfDNA were repaired and installed with the Illumina compatible adaptors. The glucosylation reactions were performed in a 25 μL solution containing 50 mM HEPES buffer (pH 8.0), 25 mM MgCl_2_, purified DNA, 100 μM N_3_-UDP-Glc, and 1 μM βGT, at 37°C for 1 hr. The reaction was purified by Micro Bio-Spin 30 Column (Bio–Rad) into ddH_2_O. After that, 1 μL DBCO-PEG4-DBCO (Click Chemistry Tools, 4.5 mM stock in DMSO) was added to the reaction mixture. The reactions were incubated at 37°C for 2 hr. Next, the DNA was purified by Micro Bio-Spin 30 Column (Bio-Rad). The purified DNA was incubated with 5 μL C1 Streptavidin beads (Life Technologies) in 2X buffer (1X buffer: 5 mM Tris pH 7.5, 0.5 mM EDTA, 1 M NaCl) for 15 min according to the manufacturer’s instruction. The beads were subsequently washed eight times for 5 min with 1X buffer. All binding and washing was done at room temperature with gentle rotation. The captured DNA fragments were amplified with 14-16 cycles of PCR amplification. The PCR products were purified using AMPure XP beads according to the manufacturer’s instructions. DNA concentration of each library was measured with a Qubit fluorometer (Life Technologies) and sequencing was performed on the Illumina Hi-Seq or NextSeq 500 platform.

#### RNA-seq library Preparation and Sequencing

Tumor and adjacent samples including two colon samples and one stomach sample were collected to isolate RNA using ZR-Duet DNA/RNA Miniprep kit (Zymo Research). Total isolated RNA was utilized to construct the library by NEBNext Ultra RNA Library Prep Kit for Illumina following the manufacture’s protocol. Sequencing reactions were executed on the NextSeq 500 platform using paired-end mode, yielding at least 32 M reads per sample.

#### PDX Preparation and Samples Collection

Establishment of patient-derived tumor xenografts: The animal protocol for this study was reviewed and approved by the Ethical Committee of Medical Research, Huashan Hospital of Fudan University. BALB/c nu/nu mice were 6–8 weeks old and weighed 16–20 g at reception (SLAC LABORATORY ANIMAL, Inc.). The fresh pathological tissue fragments were placed in sterile tissue culture medium on ice and brought immediately to the animal facility. Tumor-graft samples were cut into multiple 1x1x1 mm fragments in complete media. Tumor was implanted into female BALB/c nu/nu mice under isoflurane anesthesia, and all efforts were made to minimize suffering. A skin incision (0.3cm) was subsequently made on the right mid-back. One tumor piece (1–3mm) was inserted into each pocket and the skin was closed. Mice were regularly checked. When tumor diameter reached 1.5cm, mice were euthanized and tumors were excised, cut into 1x1x1mm fragments again, and passaged to successive generations of 3 mice. The remaining tumor was snap frozen in liquid nitrogen and stored at –80°C, and the plasma was separated from the blood sampled via the mouse eyeball. In this study, the gastric cancer and colorectal cancer patient-derived tumor xenografts were randomly selected in our existing PDX model library, while the control group was BALB/c nu/nu mice 12–14 weeks old.

#### 5hmC Enrichment Analysis

We designed two similar spike-in probes with unique sequences, named 5hmC spike-in and no5hmC spike-in. 5hmC spike-in: 5- CTGTCATGGTGACAAAGGCATCC*GGCAGAAATGCCCACACAGCCTCTTTAACCAGC ACGCCAACCGCCTCTGCTTCGGCCCTGGTCACGCAGCTGACAAGGTCTTCATAATAGAGAAATCCTG-3’, C^*^ – 5hmC modifications. no5hmC spike-in: 5’- CTGTCATGGTGACAAAGGCATCGCAGCGAAATGCCCACACAGCCTCTTTAACCAGCACGCCAACCGCCTCTGCTTCGGCCCTGGTCACGCAGCTGACAAGGTCTTCATAATAGAGAAATCCTG-3’

These sequences cannot map to the human reference genome. Six cfDNA sequencing libraries were constructed from the same cfDNA (10 ng) sample, which were divided into control and experiment groups, each having three duplicates. 100 million copies of 5hmC and no5hmC spike-ins were then mixed with the experiment sample before library preparation. The control group did not include the 5hmC pull-down step while the experiment group included the 5hmC pull-down procedure. After sequencing, we extracted spike-in reads and calculated the enrichment ratios. The average ratio of 5hmC spike-in to no5hmC spike-in in the control group was 0.72, while the ratio in the experiment group was 40.36.

#### Technical Stability Analysis for 5hmC Seal-seq Library Preparation

The designed spike-in probes were utilized to improve the robustness and sensitivity of hmC-Seal. 20 thousand copies of 5hmC and no5hmC spike-ins were pre-mixed and then added into the same cfDNA samples before library constructions. Different spike-in samples were designed as follows: 20 ng cfDNA with 2 repeats, 10 ng cfDNA with 10 repeats, 5 ng cfDNA with 2 repeats, 2 ng cfDNA with 2 repeats, 1 ng cfDNA with 2 repeats.

#### Total 5hmC quantification in cfDNA and genomic DNA

The enzymatic digestion protocol for each genomic DNA and cfDNA sample was the same. Genomic DNA or cfDNA (all in 8 μL H2O) was first denatured by heating at 95°C for 5 min and then transferred into ice water, cooling for 2 min. After that, 1 μL of 10 × S1 nuclease buffer (30 nM CH_3_COONa, pH 4.6, 260 mM NaCl, 1 mM ZnSO_4_) and 180 units (1 μL) of S1 nuclease were added into the DNA solution. The mixture (10 μL) was then incubated at 37 °C for 4 hours. Then 34.5 μL of H_2_O, 5 μL of 10 × alkaline phosphatase buffer (50 mM Tris-HCl, 10 mM MgCl_2_, pH 9.0), 0.5 μL of alkaline phosphatase were added into the DNA digestion solution. The incubation was continued at 37 °C for an additional 4 hours.

The CE-ESI-MS experiments were carried out with CESI-8000 capillary electrophoresis (CE) system from Beckman Coulter (Brea, California, USA) coupled with a Sciex Tripel Quad 5500 Mass Spectrometer (Sciex, USA) through a modified Nanosprayed?interface. Bare fused-silica capillaries etched with a porous tip were made available by Beckman Coulter (Brea, California, USA), which could be inserted into the sheathless nanospray interface. The separation capillary was 100 cm long with an internal diameter of 30 μm and an outside diameter of 150 μm. The capillary was flushed with methanol for 10 min at 100 psi, followed by water, 0.1 M sodium hydroxide, 0.1 M hydrochloric acid and water for 10 min each at 100 psi, and finally by the background electrolyte (BGE) of 10 % acetic acid (pH 2.2) for 10 min at 100 psi before first used. The BGE was also used as conductive liquid in the conductive liquid capillary. Before each run, the conductive liquid capillary was rinsed with BGE for 5 min at 100 psi. Samples for detection were stored at 5°C in the CE system. Hydrodynamic injections were used in this study, and about 100 nL sample was injected into the separation system for each analysis. A voltage of +25 kV was applied during the separation and the current was between 3.0 to 3.2 μA. The electrospray voltage was optimized to get the best nanospray stability and efficiency and +1.7 kV was good enough for this study. The quantification calibration curves of 5’-dC, 5’-mdC and 5’- hmdC were constructed using mixture solution of their standards in different concentration. (Parameters of each calibration curve were shown in Table 3) The resulting solutions of our DNA samples were directly measured by CE-ESI-MS. The concentrations of these three nucleosides in each sample were calculated based on the calibration curves. And the 5’-mdC / dC and 5’-hmdC / (dC + 5’-mdC) results of each sample were then calculated.

#### Sequencing Data Processing and Detection of Differential Loci

Read-through sequences within raw sequencing reads were trimmed using Trimmomatic version 0.35(*35*). Low quality bases at the 5’ (Phred quality score <5) and 3’ (5bp-sliding window Phred score< 15) were also trimmed. Reads with a minimum length of 50bp were aligned to the human genome assembly GRCh37 using Bowtie2 version 2.2.6(*36*) with end-to-end alignment mode. For paired-end sequencing data, read pairs were concordantly aligned with fragment length <500bp and with up to 1 ambiguous base and four mismatched bases per 100bp length. Alignments with Mapping Quality Score (MAPQ) ≥10 were counted for overlap with genomic features using featureCounts of subread version 1.5.0-p1(*37*), without strand information. Autosomal feature counts with >10 mean counts across samples were then normalized and compared between-group using DESeq2 version 1.12.3(*38*). Since gender is not a significant covariate for both autosomal gene expression(*39*) and DNA methylation(*40*), while aging has been linked to DNA methylation(*41*), age at sample collection/surgery was included as a categorical variable (<20, 20-55, >55 yr) in the negative binomial generalized linear model implemented in DESeq2. As for the experimental batch (discovery, validation or additional validation), samples were processed <1 week by 1-3 technicians, the identity of the technician was included in the model to adjust for potential technical correlation. When comparing tumor and adjacent tissues, patient identity was nested under technician identity. A FDR(*42*) of 5% was used to identify differential 5hmC loci. For PDX mouse plasma cfDNA data, sequencing reads were trimmed and aligned to a composite assembly of mixed human and mouse (GRCm38) genome. Unique alignments (MAPQ ≥10) were separated to human and mouse reads by chromosome name.

For RNA-seq data, sequencing reads were trimmed and aligned to GRCh37 annotated with GENCODE release 19, using STAR version 2.5.1b(*43*). Unique alignments with ≥90% match over reads were summarized by featureCounts. For correlation analysis in Fig.S7a, 5hmC data were also summarized over exon regions as in RNA data. For genes having >10 mean counts across samples, log_2_ fold change between tumor and adjacent tissues were estimated by DESeq2 adjusting for patient identity.

#### Refining CpG Biomarkers and Evaluating Performance

Cancer prediction models were trained using the differential 5hmC loci detected in the discovery batch. We applied elastic net regularization on a logistic linear regression model(*44*), using the *glmnet* library in the R Statistical Package(*44*):

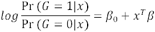

Where x is a *J*∊ (1-*j*) by *I* ∊ (1- *i*) matrix of 5hmC level at gene *j* for sample i. The model is solved by

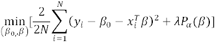

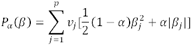

, where P _α_ is the blend of the ridge (a= 0) and the lasso (α = 1) penalty. The parameter λ controls for the overall strength of penalty, while the parameter α controls for the relative proportion between the ridge and lasso penalty. For a givenα, λ is estimated by cross-validation and selected as the largest λat which the mean cross-validated error is within one standard error of the minimum. As we expect that 5hmC loci with larger effect size are more robustly detected by the assay and therefore more reproducible across experiments, we included in the model a penalty factor *vj* so that

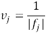

, where *tj* is the log_2_ fold change of gene *j* estimated in the training batch.

The parameter λreflects the model assumption (i.e., a large number of small effects or a small number of large effects). In small datasets like our discovery batch, the selection of α based on residual errors may lead to an over-fitted model. Instead, our model assumption was guided by the validation batch, so that α was searched to maximize AUC in the validation batch over a grid of values from 0.05 to 0.95. The model, derived from the training batch using the selected α, was applied in all classifications as described in Fig. 3 and Fig. S6.

The normalization of data in cancer classification adopts the regularized log transformation implemented in DESeq2, which estimates a global mean-dispersion trend to shrink the variance at low count genes that are associated with high Poisson noise, so that variance is stabilized across genes in the log transformed data. In Fig. 3 and Fig. S6, data from the validation batches were all normalized to the reference distribution derived from the training batch and used directly in cancer classification, i.e., we essentially ignored any remaining batch effect which could be resulted from library preparation and sequencing run. This is because that under a real clinical setting, batch effect estimated between testing samples and training samples will generally be biased, due to highly unbalanced case/control proportion in testing samples (low incidence of cancers). Batch effect may introduce some deviation from the 0.5 probability cutoff in cancer calling. External spike-in may be used to estimate batch effect in future investigations.

Receiver operating characteristic (ROC) curves(*45*) were generated to evaluate the performance of a prediction algorithm, using the *pROC*(*46*) library in the R package. Sensitivity and specificity were estimated at the score cutoff that maximizes the sum of sensitivity and specificity using the *ROCR*(*45*) library in the R package.

#### Statistical Analyses

For Fig. 4a, *P*-value was estimated by a linear mixed effects model: proportion of human reads (square root transformed) ∼ xenograft status (none | colorectal | gastric) + γ (tumor donor identity) + □ random effect γ was introduced to control for correlation among replicate xenografts. For Fig. 4b, *P*-value was estimated by a linear mixed effects model: Pearson’s *r* with tumor donor ∼ number of passages + γ (tumor donor identity) +□.

#### Annotating CpG sites with Genomic Features and Functional Analysis

The genomic features analyzed included promoters (3kb upstream of gene start), gene bodies (gene start to stop sites annotated by GENCODE release 24)(*47*), CpG islands (annotated CpG islands by UCSC Table Browser plus +/- 1kb region) and the Encyclopedia of DNA Elements (ENCODE)(*31*) features (DNase I hypersensitive sites, transcription factor binding sites, and histone modifications)(*31*). Each type of ENCODE features from the profiled ENCODE cell lines was integrated into a single list of features by collapsing overlapped and nearby (<150bp) peaks. The ENCODE features analyzed in the 5hmC genomic distribution (Fig. S2, S3) were as originally annotated without collapsing, with 20,000 features randomly sampled in the interquartile size distribution for each feature category. Pathway enrichment analysis of genes with cancer patient-associated 5hmC loci were explored based on the Kyoto Encyclopedia of Genes and Genomes (KEGG)(*48*) using the NIH/DAVID tool(*49*).

## Supplementary Materials

Fig. S1. Technical validation of the modified hmC-Seal assay using spike-in.

Fig. S2. Global 5hmC levels in plasma cfDNA and tissue gDNA.

Fig. S3. Genomic distribution of 5hmC detected in plasma cfDNA and tissue gDNA.

Fig. S4. The median distribution of 5hmC is similar between cancer and control.

Fig. S5. Differential 5hmC loci detected in cancer plasma cfDNA and tumor gDNA.

Fig. S6. Performance of 5hmC biomarkers for gastric cancer patients.

Fig. S7. Tumor associated 5hmC changes in gene regulation.

Table S1. Clinical characteristics of colorectal and gastric cancer patients and healthy controls.

Table S2. General characteristics of hepatocellular carcinoma, pancreatic cancer, thyroid cancer, gastric benign diseases, colorectal benign diseases, US colorectal cancer patients, and US healthy controls.

Table S3. Summary of samples used in 5hmC profiling.

Table S4. Summary of differential 5hmC loci in colorectal and gastric cancer detected for each feature type.

Table S5. Differential 5hmC loci in gene bodies detected at 5% FDR and 1.2 fold-change in the plasma cfDNA from discovery batch of colorectal cancer.

Table S6. Differential 5hmC loci in gene bodies detected at 5% FDR and 1.2 fold-change in the tumor gDNA from discovery batch of colorectal cancer.

Table S7. Differential 5hmC loci in gene bodies detected at 5% FDR and 1.2 fold-change in the plasma cfDNA from discovery batch of gastricl cancer patients.

Table S8. Differential 5hmC loci in gene bodies detected at 5% FDR and 1.2 fold-change in the tumor gDNA from discovery batch of gastric cancer patients.

## Acknowledgements

This work was supported in part, by grants from the National Institutes of Health (R21CA187869, P30CA060553, R01HG006827) and the Research Special Fund for Public Welfare Industry of Health (201402001). We thank Dr. H. Pickersgill (Life Science Editors) for editing this manuscript.

## Author Contributions

Conceptualization, Y.Z., W.Z., C.H., and J.L.; Methodology, X.L., Y.S., J.N., F.Y., L.W., G.J.; Investigation, X.L., Y.S.; Formal Analysis, X.Z. and W.Z.; Resources, Z.L., J.Z., W.Z., D.X., Y.W., Y.D., S.Y., J.H., J.S., H.H., F.L., L.H., P.W., X.,Q., M.C., T.Z., Q.L., M.D., Z.L., G.C., K.M., S.A., M.B.; Writing – Original Draft, W.L., X.Z., X.L., L.Y., W.Z., and C.H.; Writing – Review & Editing, Y.Z., and J.L.; Funding Acquisition, Y.Z., W.Z., and C.H.

## Confliction of Interest Statement

The University of Chicago has filed for patent protection on the original hmC-Seal technology in 2012. C.H. was one of the inventors. X.L. and Y.S. are shareholders of a company that has licensed the technology for clinical applications.

## Data Availability

All of the raw and processed data used in this study have been uploaded to the NCBI Sequence Read Archive (SRP080977) and Gene Expression Omnibus (GSE89570) depositories. The R code related to classifier detection and modeling is available upon request.

## Supplementary Materials

**Figure S1.**
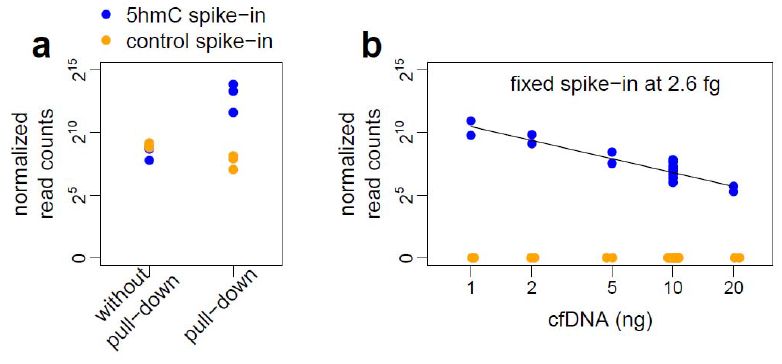
Technical validation of the modified hmC-Seal assay using spike-in probes containing 5hmC. **a**, Enrichment of 5hmC by the pull-down assay. **b**, Different amounts of cfDNA with fixed spike-in probes. The log_2_ cfDNA concentration and the mean log_2_ spike-in copy number at each concentration was close to a complete correlation (*r*^2^=0.99). Note that technical replicates, including 10 spike-in replicates with 2.6fg spike-in probes and 10 ng cfDNA performed by different individuals using different reagent batches, constituted 12% of total variance, further validated the robustness of this 5hmC-based approach using plasma cfDNA. In **a** and **b**, cfDNAs of assay samples were derived from the same biological sample. Equal copies of two spike-in probes (5hmC-containing spike-in and control non-5hmC spike-in) were added to each assay sample. cfDNA together with spike-in probes were sequenced on NextSeq 500 using paired-end 150 bp mode. The number of reads mapped to the sequence of the 5hmC- containing spike-in probe (blue) and control non-5hmC spike-in probe (orange) were counted.

**Figure S2.**
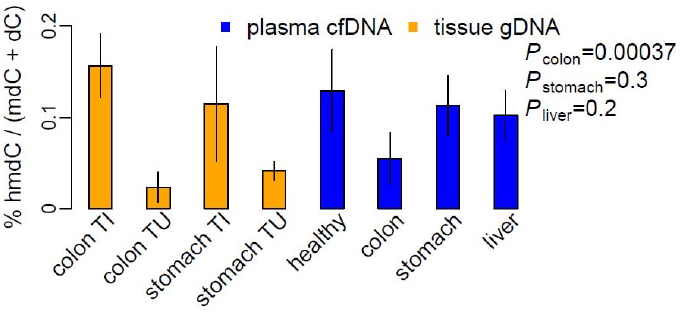
Global 5hmC levels in plasma cfDNA and tissue gDNA. P values were estimated for percent hmdC / (mdC+dC) in cancer cfDNA versus control cfDNA, by a linear model: % hmdC / (mdC+dC) ∼ cancer type (none | colorectal | gastric | liver) + age + gender + cfDNA concentration + experimental batch +□. TI: tumor adjacent tissue; TU: tumor.

**Figure S3.**
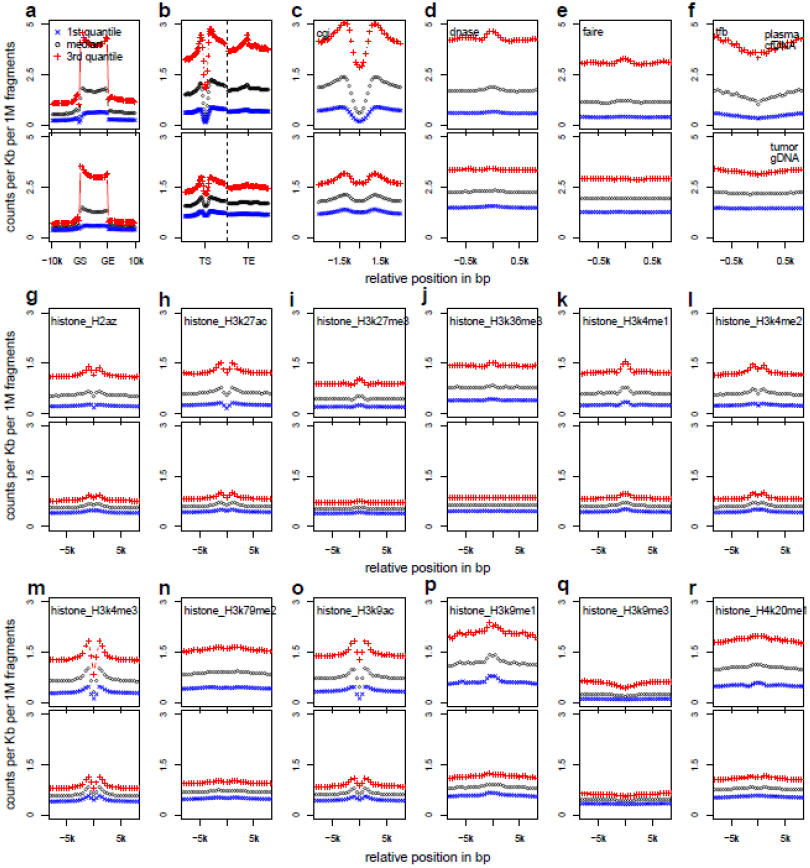
Genomic distribution of 5hmC detected in plasma cfDNA and tissue gDNA. The pooled results from the discovery batch of colorectal and gastric cancer patients as well as healthy controls are analyzed and shown. (**a-r**) shows the distribution of each feature category, respectively, for plasma cfDNA samples (upper panels) and tissue gDNA samples (lower panels). **a**, Gene bodies, defined by gene start (GS) and end (GE) sites, were divided to 20 positional bins; **b**, Regions flanking transcription start (TS) or end (TE) sites; **c**, CpG islands; **d**, DNase I hypersensitivity peaks; **e,** Formaldehyde-assisted isolation of regulatory elements; **f,** Transcription factor binding peaks; **g,** H2A.Z variant; **h**, H3K27ac; **i**, H3K27me3; **j**, H3K36me3; **k**, H3K4me1; **l**, H3K4me2; **m**, H3K4me3; **n**, H3K79me2; **o**, H3K9ac; **p**,H3K9me1; **q**, H3K9me3; and **r**, H4K20me1. In **c-r**, equal-sized bins were centered at the feature center and extended to up- and down-stream.

**Figure S4.**
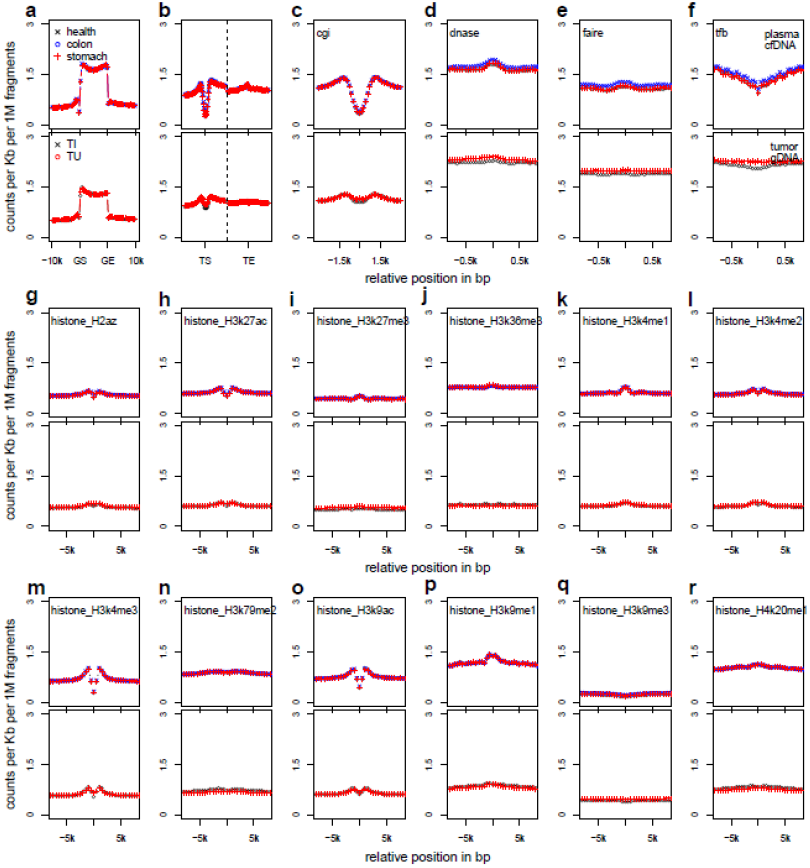
The median distribution of 5hmC is similar between cancer and control. (**a-q**) shows the distribution of each feature category, respectively, for plasma cfDNA samples (upper panels) and tissue gDNA samples (lower panels). **a**, Gene bodies, defined by gene start (GS) and end (GE) sites, were divided by 20 positional bins; **b**, Regions flanking transcription start (TS) or end (TE) sites; **c**, CpG islands; **d**, DNase I hypersensitivity regions; **e**, Formaldehyde-assisted isolation of regulatory elements; **f**, Transcription factor binding peaks; **g**, H2A.Z variant; **h**, H3K27ac; **i**, H3K27me3; **j**, H3K36me3; **k**, H3K4me1; **l**, H3K4me2; **m**, H3K4me3; **n**, H3K79me2; **o**, H3K9ac; **p**, H3K9me1; **q**, H3K9me3; and **r**, H4K20me1. In **c-r**, equal-sized bins were centered at the feature center and extended to up- and down-stream.

**Figure S5.**
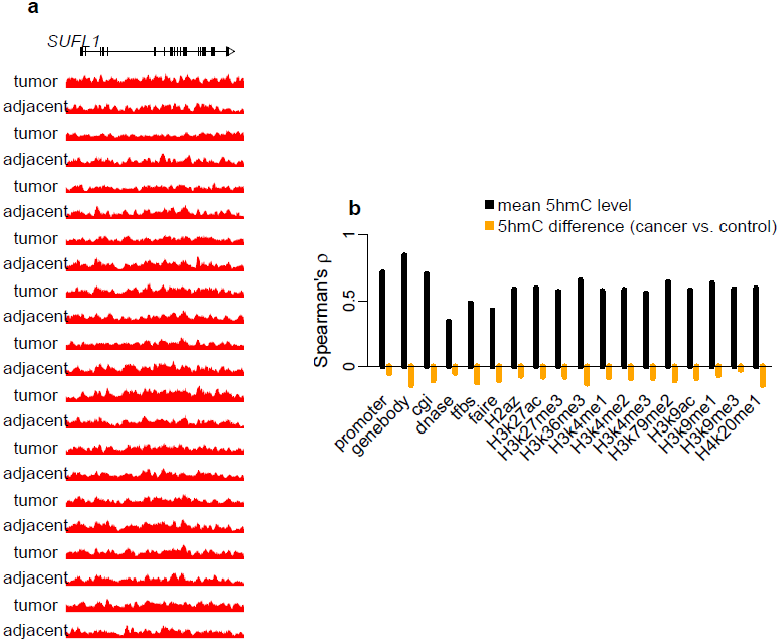
Differential 5hmC loci detected in cancer plasma cfDNA and tumor gDNA. **a**, Counts per million reads at *SULF1* gene (plus +/-20kb region) in tissue gDNA of 11 colorectal cancer patients (subset of Fig 2b). There is no significant 5hmC level difference at *SULF1* between tumor and adjacent tissues. The moving averages at 0.01 smoother span were shown. **b**, Cancer plasma cfDNA and tumor gDNA exhibit correlation in the average 5hmC levels (library-size and feature length normalized log_2_ counts, black bars), while no correlation was found for the log_2_ fold change at differential 5hmC loci detected from cancer plasma cfDNA and from tumor.

**Figure S6.**
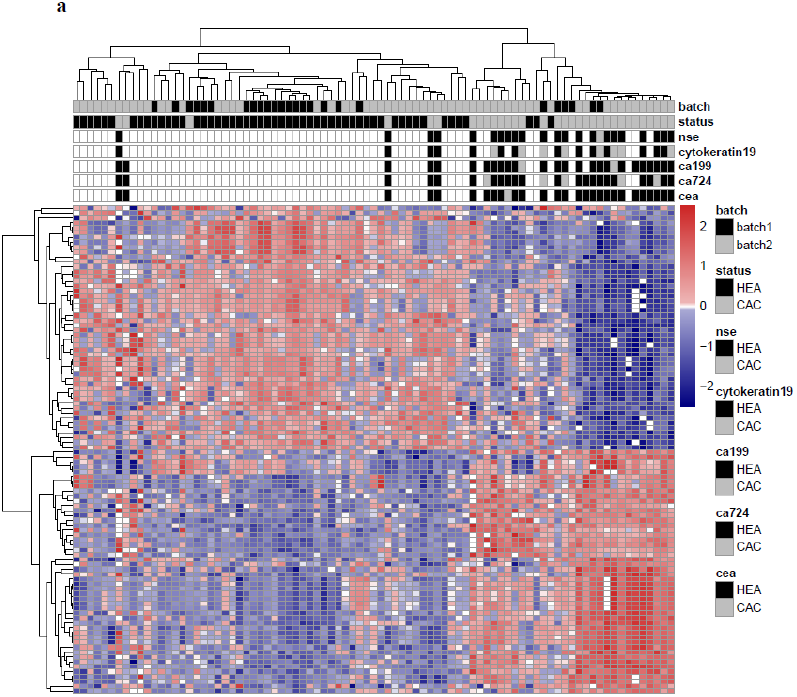
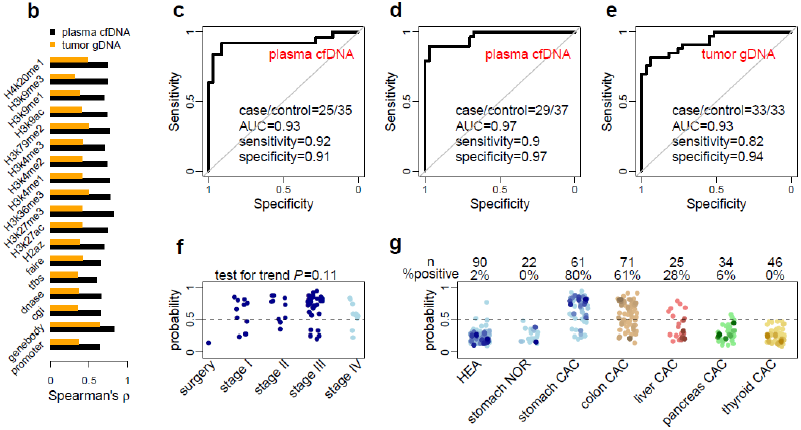
Performance of 5hmC biomarkers for gastric cancer patients. **a**, The heatmap shows clustering of cfDNA samples from both the discovery and validation batches, using the 50 most up-regulated and 50 most down-regulated 5hmC loci detected in plasma cfDNA from the discovery batch. Diagnostic results using classical biomarkers are also shown. HEA: healthy individuals; CAC: cancer patients; nse: neuron specific enolase; CEA: carcinoembryonic antigen; CA19-9/72-4: carbohydrate antigen 19-9/72-4. **b**, Correlation of 5hmC variation in cancer between the discovery and validation batches of samples is higher in plasma cfDNA (cancer patients vs. healthy individuals) than in tumor genomic DNA (tumors vs. adjacent tissues), especially for 5hmC loci in gene bodies. **c**,**d**, Classifying two independent validation batches using 5hmC classifier derived from plasma cfDNA from the discovery batch. **e**, Classifying an independent set of gastric cancer tumor tissues using 5hmC biomarkers detected from the discovery batch of tissue samples (tumors vs. adjacent tissues). AUC: area under curve. **f**, The predicted cancer probability (score) based on the 5hmC classifier from plasma cfDNA shows a trend associated with clinical stage. The one patient after chemotherapy shows a predicted probability undistinguishable from healthy individuals. **g**, The 5hmC cfDNA classifier for gastric cancer is disease- and potentially cancer type-specific, showing decreasing predicted probability in cfDNA from colorectal, liver, pancreatic and thyroid cancer patients. HEA: healthy control; NOR: patient with benign disease; CAC: cancer patient.

**Figure S7.**
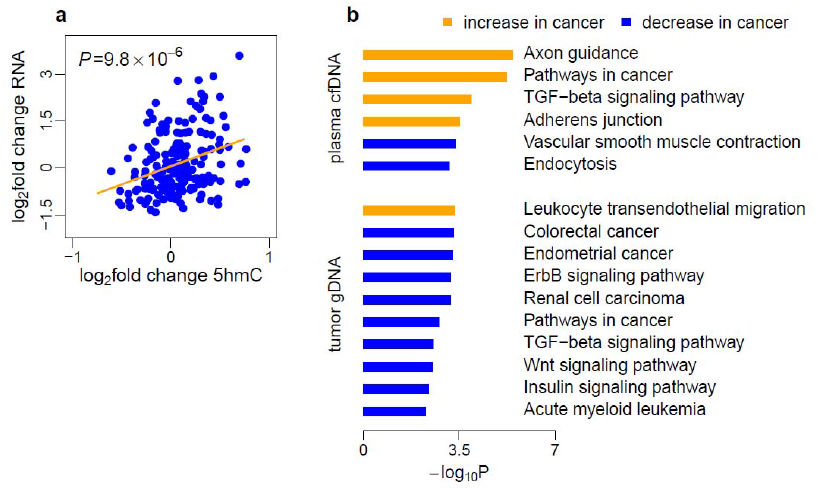
Tumor associated 5hmC changes in gene regulation. **a**, Three pairs of tumors (2 colorectal and 1 gastric) and adjacent tissues were profiled by RNA-Seq. The log2 fold changes of gene expression in tumor versus adjacent tissues were compared to the log2 fold change of 5hmC in tumor versus adjacent tissues estimated for these three patients. Gene expression changes in the tumor is positively associated with 5hmC changes in the tumor for 200 differential 5hmC loci at gene bodies detected in either colon or stomach tumors in the discovery batches. **b**, Gene pathway analysis of colorectal cancer-related canonical pathways.

**Table S1.**
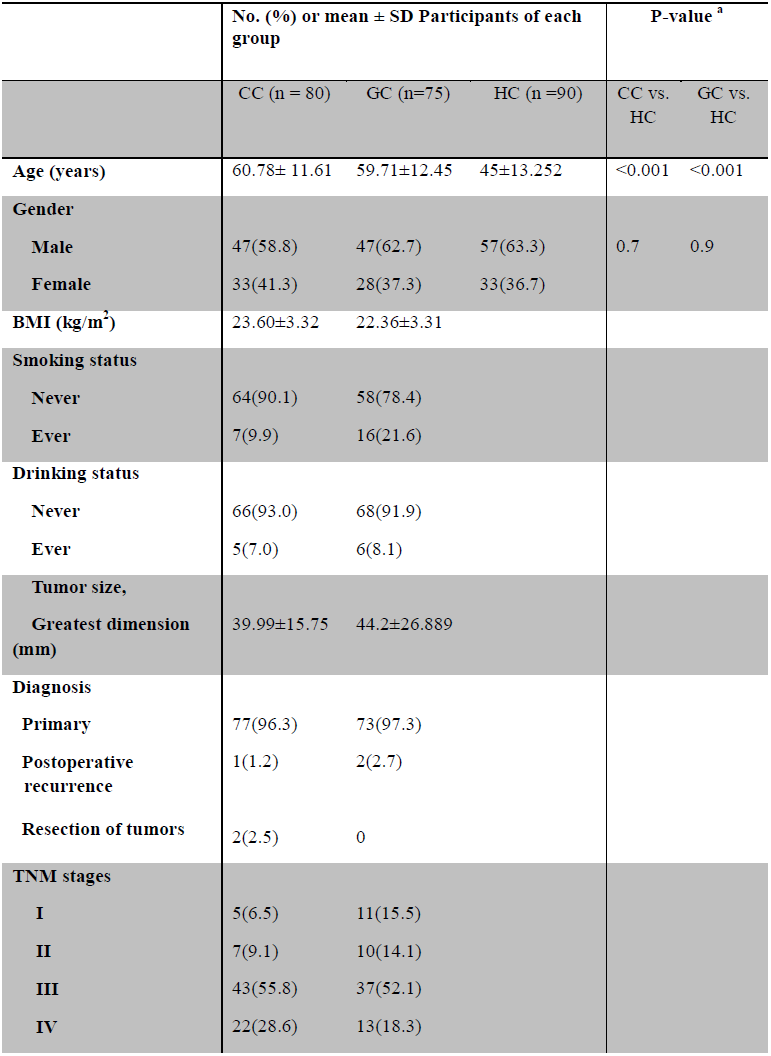
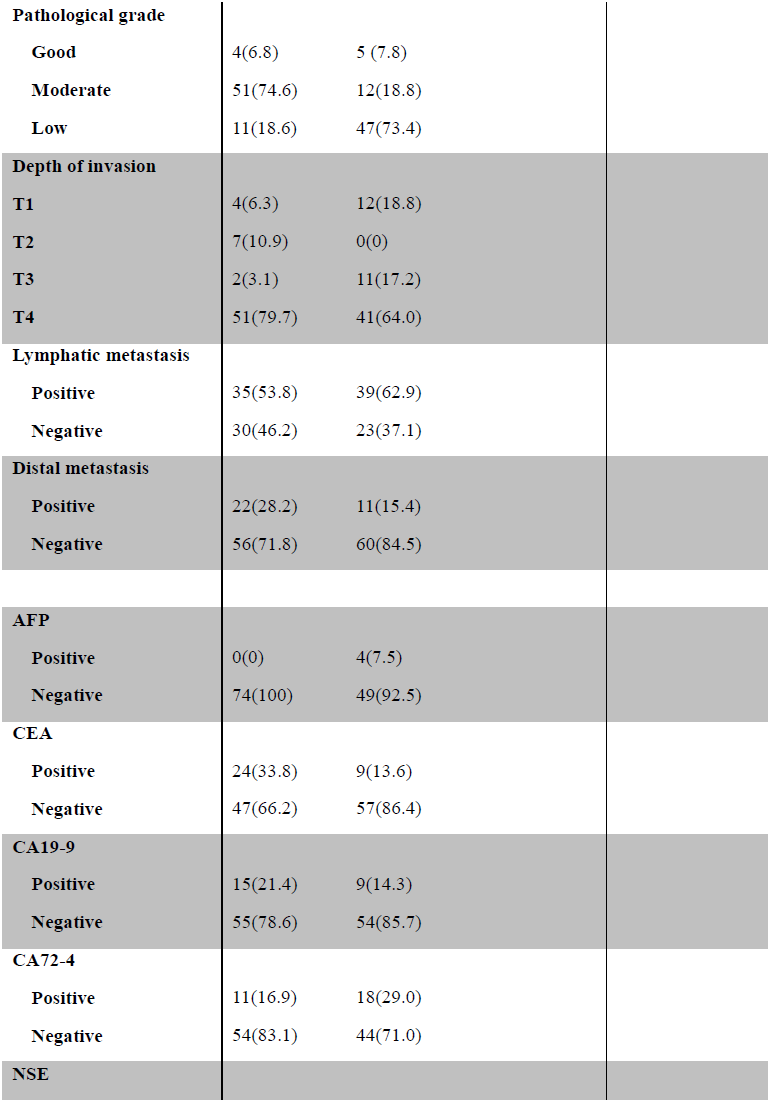
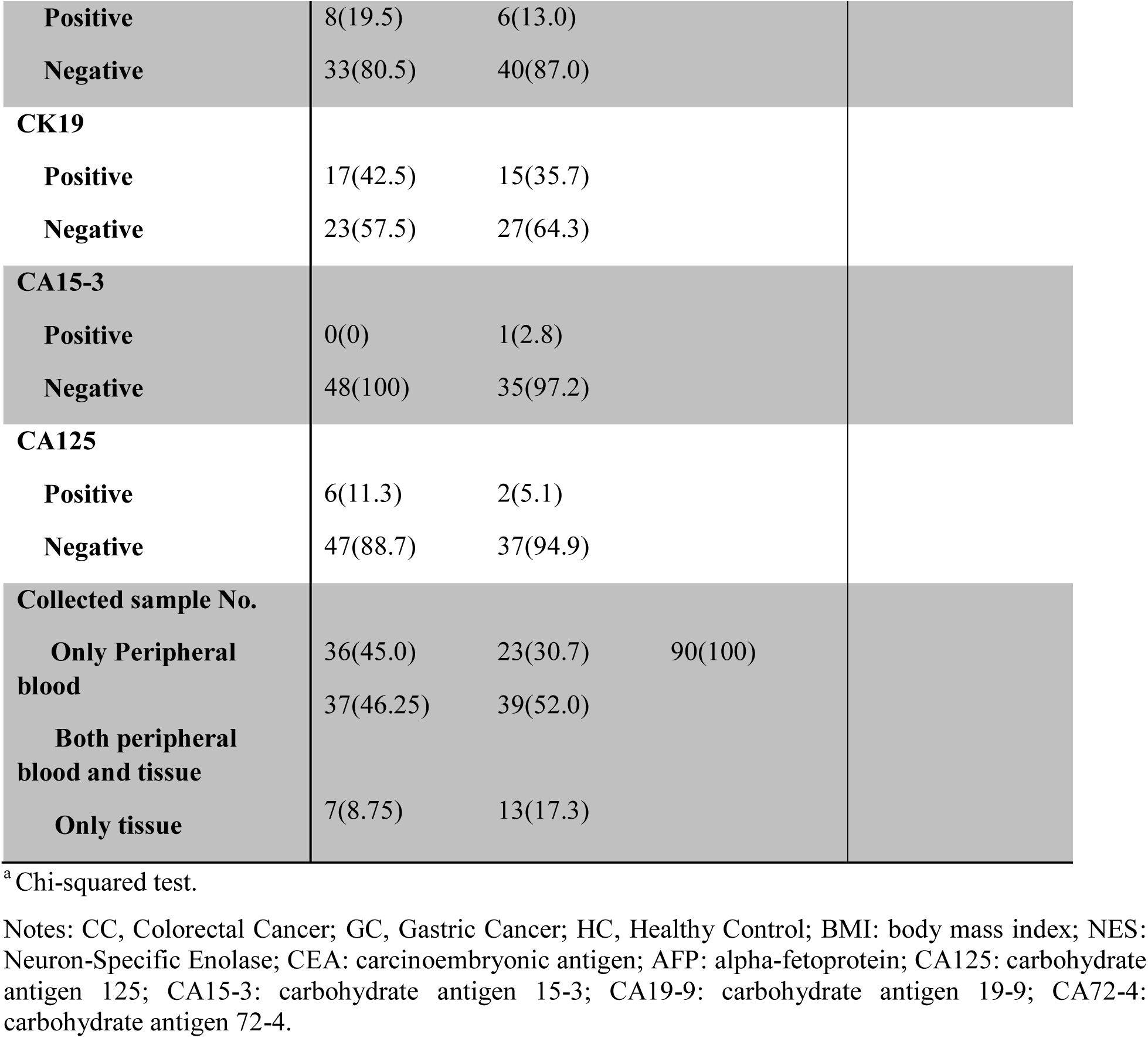
Clinical characteristics of colorectal and gastric cancer patients and healthy controls.

|  | **No. (%) or mean ± SD Participants of each group** |  |  | **P-value <sup>a</sup>** |  |
| --- | --- | --- | --- | --- | --- |
|  | CC (n = 80) | GC (n=75) | HC (n =90) | CC vs. HC | GC vs. HC |
| **Age (years)** | 60.78± 11.61 | 59.71±12.45 | 45±13.252 | <0.001 | <0.001 |
| **Gender** |  |  |  |  |  |
| **Male** | 47(58.8) | 47(62.7) | 57(63.3) | 0.7 | 0.9 |
| **Female** | 33(41.3) | 28(37.3) | 33(36.7) |  |  |
| **BMI (kg/m<sup>2</sup>)** | 23.60±3.32 | 22.36±3.31 | | | |
| **Smoking status** |  |  |  |  |  |
| **Never** | 64(90.1) | 58(78.4) |  |  |  |
| **Ever** | 7(9.9) | 16(21.6) |  |  |  |
| **Drinking status** |  |  |  |  |  |
| **Never** | 66(93.0) | 68(91.9) |  |  |  |
| **Ever** | 5(7.0) | 6(8.1) |  |  |  |
| **Tumor size, Greatest dimension (mm)** | 39.99±15.75 | 44.2±26.889 | | | |
| **Diagnosis** |  |  |  |  |  |
| **Primary** | 77(96.3) | 73(97.3) |  |  |  |
| **Postoperative recurrence** | 1(1.2) | 2(2.7) |  |  |  |
| **Resection of tumors** | 2(2.5) | 0 |  |  |  |
| **TNM stages** |  |  |  |  |  |
| **I** | 5(6.5) | 11(15.5) |  |  |  |
| **II** | 7(9.1) | 10(14.1) |  |  |  |
| **III** | 43(55.8) | 37(52.1) |  |  |  |
| **IV** | 22(28.6) | 13(18.3) |  |  |  |

|  |  |  |
| --- | --- | --- |
| **Pathological grade** |  |  |
| **Good** | 4(6.8) | 5 (7.8) |
| **Moderate** | 51(74.6) | 12(18.8) |
| **Low** | 11(18.6) | 47(73.4) |
| **Depth of invasion** |  |  |
| **T1** | 4(6.3) | 12(18.8) |
| **T2** | 7(10.9) | 0(0) |
| **T3** | 2(3.1) | 11(17.2) |
| **T4** | 51(79.7) | 41(64.0) |
| **Lymphatic metastasis** |  |  |
| **Positive** | 35(53.8) | 39(62.9) |
| **Negative** | 30(46.2) | 23(37.1) |
| **Distal metastasis** |  |  |
| **Positive** | 22(28.2) | 11(15.4) |
| **Negative** | 56(71.8) | 60(84.5) |
| **AFP** |  |  |
| **Positive** | 0(0) | 4(7.5) |
| **Negative** | 74(100) | 49(92.5) |
| **CEA** |  |  |
| **Positive** | 24(33.8) | 9(13.6) |
| **Negative** | 47(66.2) | 57(86.4) |
| **CA19-9** |  |  |
| **Positive** | 15(21.4) | 9(14.3) |
| **Negative** | 55(78.6) | 54(85.7) |
| **CA72-4** |  |  |
| **Positive** | 11(16.9) | 18(29.0) |
| **Negative** | 54(83.1) | 44(71.0) |
| **NSE** |  |  |

|  |  |  |  |
| --- | --- | --- | --- |
| **Positive** | 8(19.5) | 6(13.0) |  |
| **Negative** | 33(80.5) | 40(87.0) |  |
| **CK19** |  |  |  |
| **Positive** | 17(42.5) | 15(35.7) |  |
| **Negative** | 23(57.5) | 27(64.3) |  |
| **CA15-3** |  |  |  |
| **Positive** | 0(0) | 1(2.8) |  |
| **Negative** | 48(100) | 35(97.2) |  |
| **CA125** |  |  |  |
| **Positive** | 6(11.3) | 2(5.1) |  |
| **Negative** | 47(88.7) | 37(94.9) |  |
| **Collected sample No.** |  |  |  |
| **Only Peripheral blood** | 36(45.0) | 23(30.7) | 90(100) |
| **Both peripheral blood and tissue** | 37(46.25) | 39(52.0) |  |
| **Only tissue** | 7(8.75) | 13(17.3) |  |

<sup>a</sup> Chi-squared test.
Notes: CC, Colorectal Cancer; GC, Gastric Cancer; HC, Healthy Control; BMI: body mass index; NES: Neuron-Specific Enolase; CEA: carcinoembryonic antigen; AFP: alpha-fetoprotein; CA125: carbohydrate antigen 125; CA15-3: carbohydrate antigen 15-3; CA19-9: carbohydrate antigen 19-9; CA72-4: carbohydrate antigen 72-4.

**Table S2.**
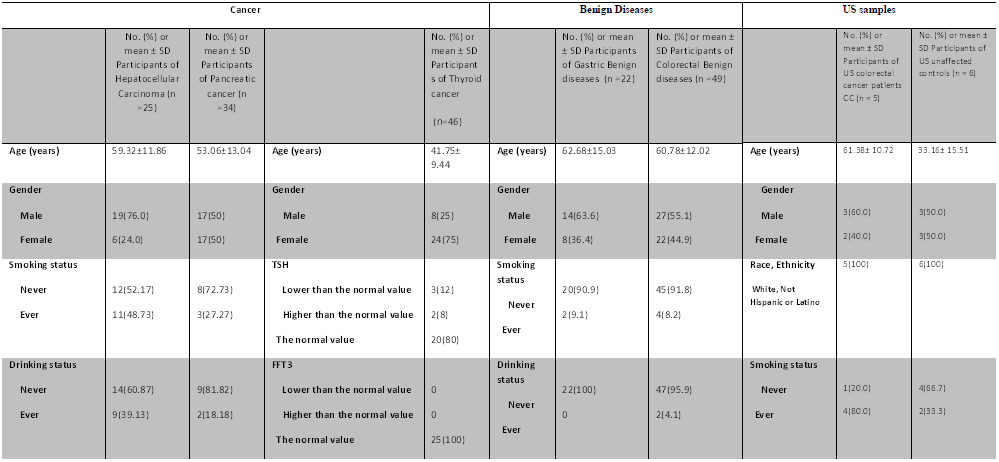
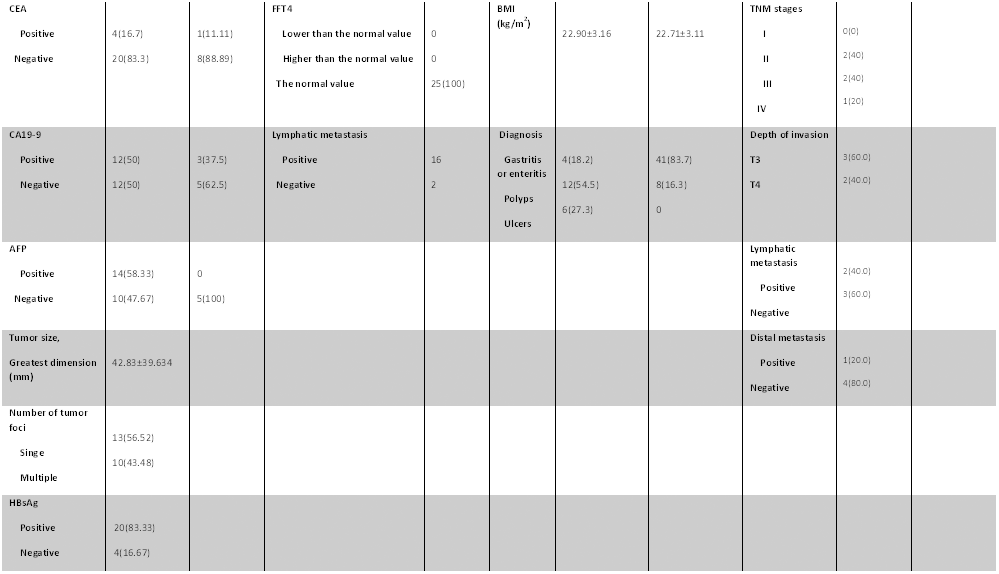
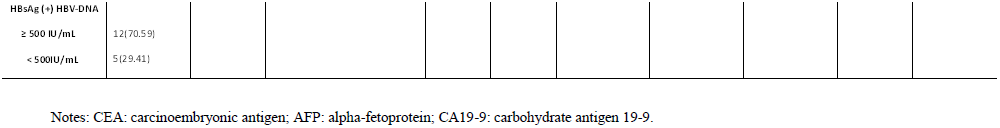
General characteristics of hepatocellular carcinoma, pancreatic cancer, thyroid cancer, gastric benign diseases, colorectal benign diseases, US colorectal cancer patients, and US healthy controls.

